# Long-distance dispersal of oilseed rape seeds: The role of grain trailers

**DOI:** 10.1101/2020.06.08.139857

**Authors:** Diane Bailleul, Sébastien Ollier, Jane Lecomte

## Abstract

In agroecosystems, anthropogenic activities can modify the natural dispersal capacity of crops and their capacity to establish feral populations. In the case of oilseed rape (OSR), seed spillage from grain trailers during harvest was first quantified by an *in situ* scientific study (Selommes, Loir-et-Cher, France). Demographic analysis of seeds collected from 85 traps set on road verges suggested that OSR dispersal distance due to seed spillage from grain trailers can be up to 400m. In the present study, we used SSR markers to genotype seeds collected from trap-sites and from surrounding OSR fields to precisely estimate the distances between traps and fields. Trailer directions on each road were also considered. Few seeds (5.8%) were not linked to a field in the studied area, while most of the seeds (59.2%) were linked to a field situated over 400 m away. The overall mean dispersal distance was 1250 m. It ranged from 308 m to 1392 m for one-lane roads, and from 1048 m to 1404 m for two-lane roads. Events of seed dispersal at greater distances (> 5 km) were rare but still possible. It thus follows that OSR seed dispersal due to spillage from grain trailers should be carefully considered in the context of genetically modified plant cultivation.

## Introduction

Seed dispersal is often a complex phenomenon that governs the dispersal of annual plants [1]. It is crucial to quantify this dispersal in order to understand the population dynamics and thus the spatial distribution of the species [2]. In areas where human activities are intense, human-mediated seed dispersal (i.e., anthropochory) considerably affects plant dispersal [3,4].

Different anthropogenic vectors can disperse seeds: humans [5,6], cars [7-12], and agricultural machinery in agroecosystems. Whereas cars and humans are able to disperse limited quantities of seeds but across large distances [3,8,13], agricultural machinery can disperse large amounts of seeds. In this case, seeds can be scattered by mowing machinery [14,15], harvesters [16,17], and grain trailers [18] either in situ or while being driven [16,18,19]. To be able to characterize seed dispersal [2], only a few rare studies have been conducted to both measure the amount of seed dispersed and the dispersal distances [15].

In agroecosystems, seed dispersal could lead to feral crop populations being established in uncultivated areas such as road verges. This is the case for oilseed rape (OSR, *Brassica napus* L.), which is one of the most commonly cultivated crops in the European Union (6 877 000 ha cultivated in Europe in 2018; France: 23.5% of the total European Union harvested area of oilseed [20]). Feral OSR populations are frequent along roadsides [16,21-25] and railways [26-28]. This is partly due to OSR pod shattering (8,000 seeds per m2 [29]; 9 to 56 times the number of seeds sown [30]), while OSR seeds also have the ability to establish long-lived seed banks in the soil for up to 17 years [31-34] via secondary dormancy depending on the type of cultivar and cultivation conditions [35-37].

In the case of genetically modified (GM) cultivation, it has been shown that feral OSR populations could arise from the spillage of imported seeds during transportation [16,38-41] or the cultivation of GM OSR populations [23,42], thus even after GM cultivars are no longer used [43,44]. Feral GM OSR populations are able to exchange genes through pollen flow with other feral GM OSR populations as well as GM OSR fields [22,23,45]. As GM OSR seeds also exhibit secondary dormancy, their survival in feral soil seedbanks could constitute reservoirs of transgenes [25] and thus increase the persistence of transgenes in the environment, which can lead to ecological, agricultural, and economic problems. The GM traits nowadays on the market for OSR plants have no reason to affect their seed dispersal abilities, thus studying non-GM OSR plants would lead to the same results and conclusion as on GM OSR plants.

The origin of feral OSR populations seems to be linked to anthropochory. They could originate from feral seed banks [24], the harvesting of adjacent fields [24], truck spillage [21,38,46], grain trailers [18], trains [47], and vehicular transport [48-50]. Therefore, the characterization of OSR seed dispersal is necessary in order to estimate the risk of long-distance seed dispersal from cultivated fields and, consequently, the establishment of feral populations. von der Lippe and Kowarik [48,49,51] quantified the dispersal of OSR seeds by vehicles using seed trapping experiments. A seed deposition experiment conducted by Garnier and Lecomte [52] showed that secondary dispersal was correlated with traffic intensity, was local, and did not systematically occur along road verges. These studies were only a quantification and not a characterization of seed dispersal, because the collected seeds could not be related to the fields of their study sites.

Bailleul *et al.* [18] performed a scientific study using OSR traps along road verges in an agricultural landscape. They laid seed traps on road verges located near OSR fields (at a distance of 0 m, 40 m, and 400 m from the OSR fields) to measure seed spillage from grain trailers during harvest. They used a statistical model to explain the amount of trapped seeds and found that the number of seeds in traps depended on the trap-field distance as well as the distance to the main silo in interaction with the number of road lanes. However, this approach did not permit an accurate conclusion to be drawn in terms of an estimate of the effective dispersal distances of the seeds. For example, a seed found in a trap placed next to an OSR field, that is to say, at a distance of 0 m, could come from this field or indeed another field situated further away.

Based on this scientific study and the analysis of the seeds collected in the traps and fields in the area, we used microsatellite markers to estimate precise seed dispersal distance due to seed spillage from grain trailers. We discuss these results in the framework of GM cultivation and the coexistence of GM, non-GM, and organic crop cultivation.

## Materials & Methods

### Study area

The study area is a typical open-field agricultural landscape of 41km2, centered around the village of Selommes (Loir-et-Cher, France, 47°45’24’’N; 1°11’34’’E), which has a grain silo where most local farmers take their harvested grains (see Bailleul *et al.* [18] and Fig 1). Grain trailers are used to transport seeds from harvested fields to this silo. In 2010, 118 fields of winter OSR were cultivated in this area over a total of 684 ha (16.7% of the area). Feral OSR populations were found on 10–14% of the road verges, and this proportion was the same for the last ten years [53]. Fields, feral populations, roads, and many landscape elements were recorded and mapped. Roads were categorized as paths (42% of roads) and one- (29%) or two-lane (29%) paved roads.

**Fig 1:**
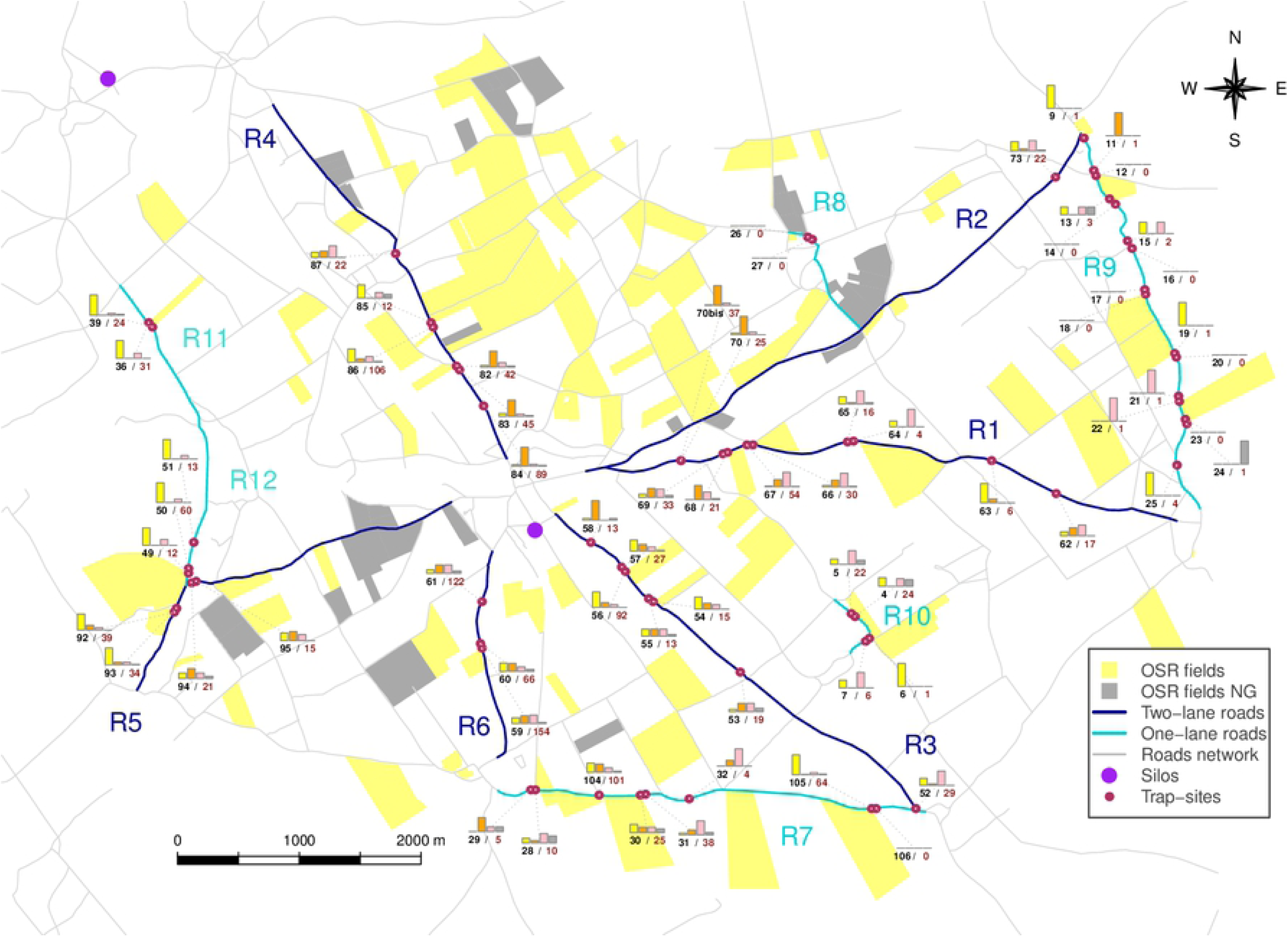
Global map of the Selommes area with origin pattern barplots of OSR seed collected at each trap-site. Seed amounts are percentage transformed; all barplot axes have the same scale between 0 and 1 (which indicates 0% to 100%). The yellow barplot corresponds to neighboring field origin, orange barplot to fields along the grain trailer trips, and pink barplot to fields elsewhere in the study area. The gray barplot corresponds to no field in the area. Each trap is associated with a number (black) and amount of seeds trapped (brown). Each road is represented by an R-number code.

No specific permission was required for this study, as French roads are public areas. Road verges are not privately owned or protected. The DDE (Direction Départementale de l’Equipement, in charge of road verge management), town mayors, and farmers were informed about the study before the experiment. The field studies did not involve endangered or protected species.

### Study design

In 2010, 85 traps at all were set (at 0 m, 40 m, and 400 m from adjacent fields, Fig 1) on six one-lane and six two-lane roads of more than 600 m in length (details on Tab 1). We used the same reference as Bailleul *et al.* [18] to characterize grain trailer trips between fields and the main silo. These traps were placed at the beginning of the OSR harvest season in 2010. Over the following eight days, all traps were checked daily, and trapped seeds were collected. At the end of the experiment, all the surrounding fields had been harvested. Grain trailers were the only direct seed transportation vehicles that circulated during harvest (note, however, the possible secondary transportation by cars).

**Tab 1:** Trap-sites information. For each trap-site, this table indicated its unique identification number (ID_site), the identification of the road (ID_road) on which the trap is set, the number of lanes (Lanes) of the road, the number of seeds trapped during the study and the distance between the trap and the nearest field (Dist_field) on the same side of the road.

| ID_site | ID_road | Lanes | Seeds_trapped | Dist_field |
| --- | --- | --- | --- | --- |
| 62 | R1 | 2 | 24 | 400 |
| 63 | R1 | 2 | 14 | 400 |
| 64 | R1 | 2 | 28 | 0 |
| 65 | R1 | 2 | 25 | 40 |
| 66 | R1 | 2 | 40 | 0 |
| 67 | R1 | 2 | 60 | 40 |
| 68 | R1 | 2 | 34 | 0 |
| 69 | R1 | 2 | 71 | 40 |
| 70 | R1 | 2 | 33 | 400 |
| 70bis | R1 | 2 | 42 | 400 |
| 4 | R10 | 1 | 27 | 0 |
| 5 | R10 | 1 | 30 | 40 |
| 6 | R10 | 1 | 1 | 0 |
| 7 | R10 | 1 | 8 | 40 |
| 36 | R11 | 1 | 31 | 0 |
| 39 | R11 | 1 | 24 | 0 |
| 49 | R12 | 1 | 12 | 0 |
| 50 | R12 | 1 | 60 | 40 |
| 51 | R12 | 1 | 13 | 400 |
| 26 | R8 | 1 | 0 | 0 |
| 27 | R8 | 1 | 0 | 40 |
| 73 | R2 | 2 | 39 | 400 |
| 74 | R2 | 2 | 65 | 0 |
| 75 | R2 | 2 | 69 | 0 |
| 76 | R2 | 2 | 54 | 40 |
| 77 | R2 | 2 | 137 | 0 |
| 78 | R2 | 2 | 6 | 40 |
| 79 | R2 | 2 | 57 | 0 |
| 80 | R2 | 2 | 100 | 40 |
| 81 | R2 | 2 | 208 | 400 |
| 102 | R2 | 2 | 235 | 0 |
| 103 | R2 | 2 | 318 | 0 |
| 52 | R3 | 2 | 36 | 400 |
| 53 | R3 | 2 | 32 | 400 |
| 54 | R3 | 2 | 21 | 0 |
| 55 | R3 | 2 | 20 | 40 |
| 56 | R3 | 2 | 199 | 0 |
| 57 | R3 | 2 | 36 | 40 |
| 58 | R3 | 2 | 20 | 400 |
| 82 | R4 | 2 | 67 | 0 |
| 83 | R4 | 2 | 106 | 40 |
| 84 | R4 | 2 | 160 | 400 |
| 85 | R4 | 2 | 13 | 0 |
| 86 | R4 | 2 | 225 | 40 |
| 87 | R4 | 2 | 35 | 400 |
| 88 | R4 | 2 | 227 | 0 |
| 89 | R4 | 2 | 375 | 40 |
| 90 | R4 | 2 | 71 | 400 |
| 92 | R5 | 2 | 286 | 0 |
| 93 | R5 | 2 | 181 | 40 |
| 94 | R5 | 2 | 26 | 0 |
| 95 | R5 | 2 | 23 | 40 |
| 96 | R5 | 2 | 31 | 0 |
| 97 | R5 | 2 | 17 | 40 |
| 98 | R5 | 2 | 102 | 0 |
| 99 | R5 | 2 | 91 | 40 |
| 100 | R5 | 2 | 96 | 0 |
| 101 | R5 | 2 | 147 | 40 |
| 59 | R6 | 2 | 330 | 0 |
| 60 | R6 | 2 | 88 | 40 |
| 61 | R6 | 2 | 158 | 400 |
| 28 | R7 | 1 | 18 | 0 |
| 29 | R7 | 1 | 8 | 40 |
| 30 | R7 | 1 | 39 | 0 |
| 31 | R7 | 1 | 44 | 40 |
| 32 | R7 | 1 | 6 | 400 |
| 104 | R7 | 1 | 120 | 0 |
| 105 | R7 | 1 | 114 | 0 |
| 106 | R7 | 1 | 1 | 40 |
| 9 | R9 | 1 | 141 | 0 |
| 11 | R9 | 1 | 2 | 40 |
| 12 | R9 | 1 | 2 | 0 |
| 13 | R9 | 1 | 94 | 0 |
| 14 | R9 | 1 | 3 | 40 |
| 15 | R9 | 1 | 2 | 400 |
| 16 | R9 | 1 | 0 | 400 |
| 17 | R9 | 1 | 0 | 40 |
| 18 | R9 | 1 | 0 | 0 |
| 19 | R9 | 1 | 1 | 0 |
| 20 | R9 | 1 | 0 | 40 |
| 21 | R9 | 1 | 1 | 40 |
| 22 | R9 | 1 | 1 | 0 |
| 23 | R9 | 1 | 0 | 0 |
| 24 | R9 | 1 | 1 | 40 |
| 25 | R9 | 1 | 6 | 400 |

### Data collection

In each trap, seeds were collected daily for eight days and then grown in a greenhouse at Université Paris-Saclay (Orsay, Essonne, France). If the trap contained less than 50 successfully germinated seeds, all the plants were used for genetic analysis by taking one leaf per plant. Otherwise, only 50 randomly selected plants were used.

We genotyped 3104 leaf samples taken from the seed-traps. Several seeds failed to germinate. Every trap with less than 50 plants (except one) was genotyped. Nearly every trap with more than 50 plants (except 7 traps) were genotyped.

OSR field plants were sampled by taking one leaf per plant. Ten plants were sampled per field. We were able to genotype a total of 1015 leaf samples from 97 of the 118 fields.

For cultivar assignment, we previously had genotype information about 58 cultivars [54] that were potentially sown in the area of Selommes from 2002 to 2005: 45 pure-line cultivars (homogeneous homozygous), 11 hybrid cultivars (homogeneous heterozygous), and 2 cultivar associations (heterogeneous genotypes).

We obtained certified seeds from 20 more recent cultivars (12 pure-line and 8 hybrid cultivars) and completed them with seeds of the 50 most probable previous cultivars that were still grown. We genotyped a total of 1976 cultivar seeds with most of the time 30 seeds per cultivar.

### Molecular markers

We selected 9 SSR markers that exhibited high polymorphism and allowed the discrimination of cultivars: 7 SSR markers previously used (Ra2E11, Na12D08, Na10H03, Ol12F02-A/Ol12F02-B, Ol11B05, Na14H11, and Ra2A05) and 2 new SSR markers (Na12C08 and Na12E01-A/Na12E01-B). Primers for Ol12F02 and Na12E01 amplified two polymorphic loci each.

Thus, in total, 11 usable polymorphic and independent loci were amplified from these 9 SSR markers.

Molecular laboratory work was conducted using the Genotyping Platform of Clermont-Ferrand (INRA, France).

### Field and seed-trap plant assignment

#### Cultivar assignment

Assignment methods were used to assign a cultivar to each sampled plant based on genotype data: exact compatibility assignment and maximum likelihood. Field leaves were assigned by the direct method of exact compatibility assignment: if the plant genotype is compatible for all of the loci with one of the genotypes of a given cultivar, the plant is assigned to this cultivar. A maximum likelihood assignment method was developed for to assign a cultivar to each of the plants sampled using seeds assignment (see Supporting Information S2 in Bailleul *et al.* [54]). Cross-recombination among cultivars was not allowed and only considered within cultivars; we also did not consider inter-cultivar hybrids. If the findings were ambiguous, the plant was assigned to the most consensual cultivar with the highest likelihood.

#### Field assignment

Due to sequencing problems (see Results section), cultivar assignments were not possible. We decided to directly assign fields to each grain from the trap-sites without cultivar identification. We kept only unique and entire field genotypes. Contrary to cultivar assignment (i.e. a field is assigned to a particular cultivar if at least six of the ten sampled plants belonged to the same cultivar), field assignment could result in several field assignments, as several fields could share the same cultivars and/or genotypes. Thus, every likelihood result that differed from 0 was considered.

### Minimum distance with the information about grain trailer trips

We computed the distances between each trap and each field in our area based on the road network. We then coupled this geographic distance information to the field assignment information to extract the minimum geographic distance between each trapped seed linked to a field in the area. Taking into account the information about the most probable trip travelled by each grain trailer from a given field to the main silo, we computed the corrected minimum distance (compared to Supplementary Information S1) between a trapped seed and the most proximal field from which the seed would have originated. We then categorized these distances as seeds coming (1) from the neighboring field, (2) from a field along the grain trailer trip (but not the neighboring field), (3) from the studied area (but not from a field along the grain trailer trip), and (4) from a non-genotyped field in the area.

The data analyses and figures generation were done with R software version 3.6 [55]. The datasets generated during the current study are available in the Dryad repository [will be accessible on Dryad after journal acceptation].

## Results

### Cultivar genotypes

Due to sequencing problems on cultivars, we did not obtain any alleles from Ol11B0 and, Ol12F02-A/Ol12F02-B. The two new markers (Na12C08 and Na12E01-A/Na12E01-B) returned a missing data percentage of 38.6% (against 7.2% for the other markers). Cultivar genotypes were thus unusable.

### Data collection genotypes

The two new markers (Na12C08 and Na12E01-A/Na12E01-B) returned 53% of missing data for field and seed-trap genotypes (against 15.1% for the other markers).

We thus focused only on the former eight SSR loci for all the field and seed-trap genotypes. On this basis, for seed-trap genotypes, we only considered genotypes with a maximum of 6 missing alleles (for a total of 16 alleles). We finally obtained 2923 proper genotypes.

For field genotypes, as they constituted our references, we only considered entire genotypes with no missing data. We obtained 865 full genotypes from 98 fields (only 1 field was discarded). We then constituted a field database reference of 444 genotypes with unique genotypes for each field.

### Field and seed-trap plant assignment

For trap-site seeds, only 131 seeds were not linked to a field genotype. This means that 2792 (over 2923, i.e. 95.5%) seeds had a potential field origin in our area. From the perspective of the fields, every field was assigned to at least one trapped seed.

### Minimum distance with the information on grain trailer trips

These field assignment results combined with the geographical distances between each trap and field enabled us to extract the minimum distance between each trapped seed and corresponding field. Taking into account the trailer trips from the fields to the main silo, we corrected the initial distances (see Supplementary Information S1): each seed was linked to the closest field according to the driving direction and farmer trailer trips (Fig 1 & 2). As some putative fields were not genotyped for seeds from roads R2, R4, and R5, minimal distances were not considered from these roads. The other roads were not impacted by the missing fields.

**Fig 2:**
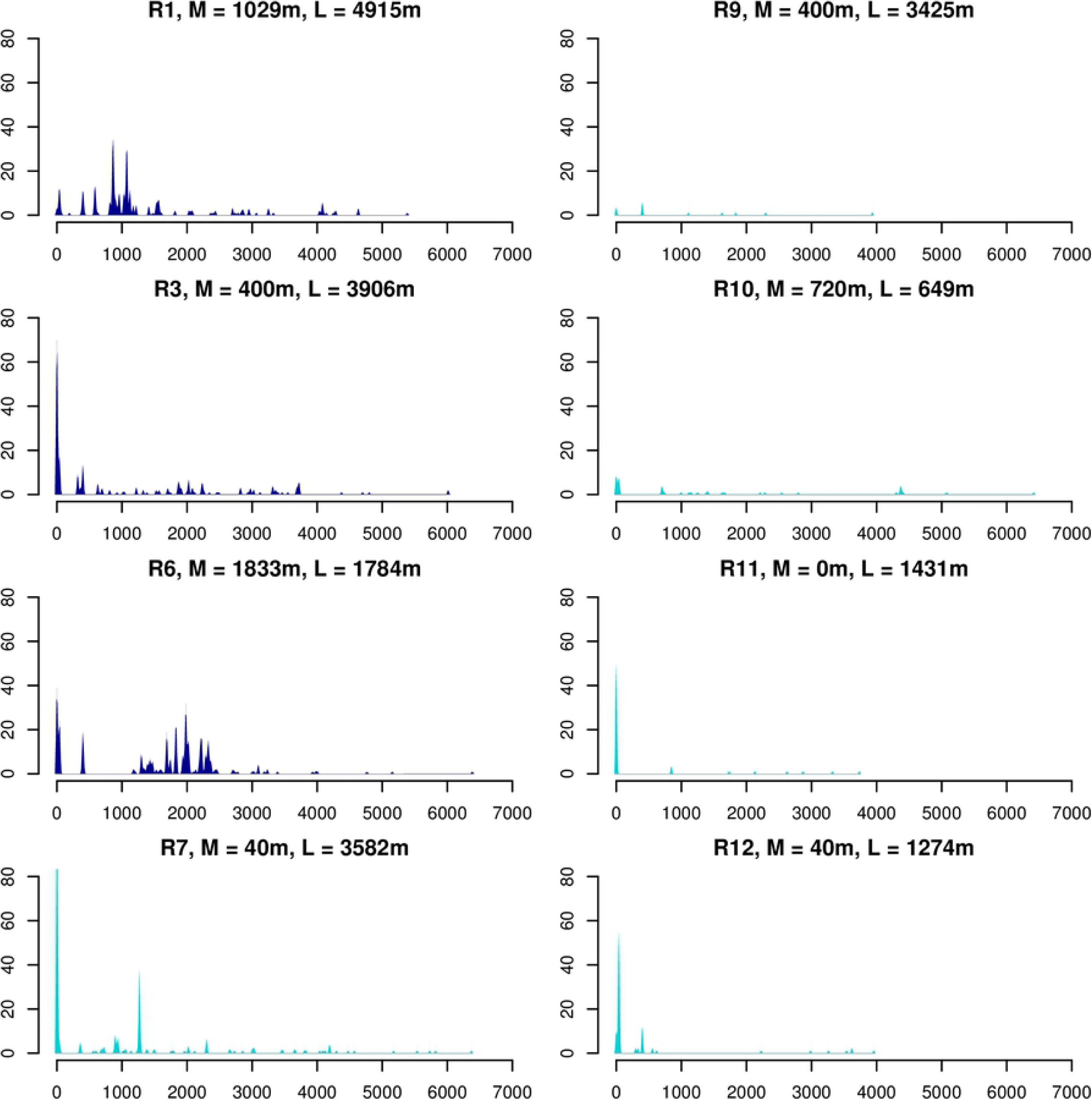
Minimal estimated distances between traps and fields. Minimal dispersal distances are on x-axis. Seeds number on y-axis. Dark blue graphs correspond to two-lane roads and light blue graphs to one-lane roads. M stands for median as “the median minimal distances between trap-sites on the road and assigned fields while taking into account grain trailer trips”. L stands for length as “total length of the road”.

For the remaining 1248 seeds, the overall mean minimal dispersal distance was 1164.7 m (median: 903 m, standard deviation -sd-: 1237.6). For two-lane roads, their mean values ranged from 1106.5 m for R3 (median: 400, sd: 1373.9, Fig 2) to 1546.7 m for R6 (median: 1833, sd: 1006.1). For one-lane roads, their mean values ranged from 351 m for R11 (median: 0 m, sd: 902.2) to 1475.6 m for R10 (median: 720 m, sd: 1788.7).

Overall these 1248 seeds, 19.6% (245) of trapped seeds were linked to a field at 0 m, 9.3% (116) to a field at 40 m, and 4.6% (58) to a field at 400 m. A huge number of seeds (688, 55.1%) was linked to a field further than 400 m away. 40% of the seeds were linked to a field further than 1165 m, the overall mean minimal dispersal distance estimated. A few seeds (12, less than 1%) were associated with very long dispersal distance (> 5 km). Additionally, 5.8% (72) were not linked to a field in this area. Some seeds (5.5%, 69) were linked to fields at distances between 0 m and 400 m.

### Categorized distances with the information on grain trailer trips

We also summarized the results with barplots for each trap with the following categories (Fig 1): seeds linked to neighboring fields, seeds linked to another field according to the grain trailer trips, seeds linked to a field elsewhere in the area, and seeds linked to no fields. The map indicates all the roads.

For all the roads (i.e., with the exception of R2, R4, and R5), 36.5% (457) of seeds were linked to their neighboring field and 27.6% (344) to a field along the grain trailer trips.

On one-lane roads, 54.9% of seeds were linked to the neighboring field (from 32.1% for R10 to 83.5% for R12), 12.5% to a field along the grain trailer trips (from 0% for R10, R11 and R12 to 22.7% for R7), and 25.3% to a field elsewhere in the area (from 0% for R12 to 20.8% for R10).

On two-lane roads, 26.1% of seeds were linked to the neighboring field (from 10.7% for R1 to 47.6% for R3), 36.2% to a field along the grain trailer trips (from 26.4% for R3 to 46.1% for R1), and 32.8% to a field elsewhere in the area (from 2.9% for R1 and R3 to 7.6% for R6).

## Discussion

The demographic analysis of the study of Bailleul *et al.* [18] suggests that OSR seed dispersal distance due to spillage from grain trailers can be up to 400 m. In this article, 400m was considered as long distance dispersal and the farther long distance dispersal ever quantified. However, our present study shows for the first time that seed dispersal can go far beyond 400 m, as it was the case for more than two-thirds of the seeds trapped on road verges. The average seed dispersal distance is 1164.7 m (or 847 m if we do not considered the information obtained from local farmers, Supplementary Information S1) and 40% of the seeds collected were linked to a field farther than this average distance. This information from grain trailer trips was thus essential to properly assign a seed to its source field. Even events of seed dispersal at “very-long” distances (> 5 km) are rare (less than 1%) but still possible. As seeds trapped on road verges were assigned to the closest and most likely field from where they could originate, we should state that our seed dispersal distances are certainly underestimated.

Our estimation of seed dispersal distance at the scale of an agricultural landscape confirms the existence of a seed flow over long distances via grain trailers. This is indeed far from the classic pattern of seed loss mostly attributed to the edge of the harvested field, because this represents only one-third of the trapped seeds. Given the considerable density of seeds dispersed (400 seeds per m2 [18]) and their viability (77%), the establishment of feral populations at a relatively long distance from the source field seems more than likely.

The knowledge of grain trailer trips was crucial in this study. It was improbable that grain trailers did not take the most direct road leading to the silo. However, in two cases, the large dispersal distance we observed could be due either to the fact that the trailers take an alternative route or that seeds experiencing a rare and intense dispersal event. As the number of collected seeds is related to the distance between a trap and the road verge, we thought that a trap placed on a side of the road could not collect seeds from a field located on the opposite side. Nevertheless, we found one case (site 29) on the narrow one-lane road R7 where three seeds seemed to originate from the field on the other side of the road.

From the genetic analysis of this study area, we showed that the diversity of the feral population (i.e., successful seed dispersal and survival) in terms of cultivars was greater on road verges than on path verges [54]. We thus hypothesized that the more intense the traffic on a road, the more seeds from different fields were spilled by grain trailers. This issue was difficult to address in this study, as we could not use cultivars as a proxy. However, we showed that on average, more seeds were linked to neighboring fields on one-lane roads than two-lane roads, thus suggesting that dispersal is greater on two-lane roads. Grain trailers are likely to drive faster on two-lane roads, and as a result, seed dispersal due to wind blowing would be greater.

These results should be interpreted with caution, as a number of choices limit their scope. In terms of assignment, we retained the missing data for some seed genotypes, which limits the reliability of the procedure. However, the genotypes with missing data are relatively few. Also, only 10 leaves per field were genotyped, which is likely to limit the characterization of the genetic diversity of the seed sources. However, we were able to show that 10 leaves allow a non-negligible part of this diversity to be estimated for a reasonable sampling effort and genotyping cost (in Supporting Information [54]). Finally, we chose to assign the seeds to the nearest field whose likelihood was non-zero, which leads to an underestimation of the dispersal distance.

This study faced some limitations. First, an inherent difficulty in this study relates to the very nature of the organization of biodiversity in the agroecosystem under investigation. As the diversity of OSR cultivars is relatively low, and these cultivars are relatively homogeneous in term of genotypes, it is therefore difficult to discriminate them with only a few genetic markers. Second, the fact that some fields could not be genotyped forced us to disregard three roads, which would have had an impact on the proportion of seeds linked to another location in the study area. Another potential bias could potentially originate from the fact that the frequency estimation of seed origin could differ between trap were all the plants were analyzed (trap with less than 50 seeds) and those were 50 seeds were randomly selected.

### Perspectives

To provide a more accurate estimate and be able to disentangle the processes involved in seed dispersal due spillage from grain trailers at the agroecosystem scale, an experiment should be conducted in collaboration with farmers in order to better control the sources of seed dispersal and their location in space, while using sufficiently different cultivars in term of genotypes. In the context of GM plant cultivation, the long seed dispersal distance of GM OSR seeds that we found in this study should be taken into account in scenarios of coexistence between GM, non-GM, and organic OSR. Because OSR seed is small and light, appropriate management policies such as covering the top of grain trailers or not overfilling trailers will be challenging to implement due to the economic constraints on farmers. Indeed, not filling trailers to capacity means more trips and thus higher fuel consumption. Moreover, covering the trailers with a tarp increases the risk of seed decommissioning when they are marketed at the silo.

## Acknowledgements

We thank Fanny Loesch for the field support and Clarisse Pelisse for the greenhouse work with the help of Lionel Saunois and Amandine Dubois. We also thanks Agnès Ricroch for the obtainment of certified seeds and farmer surveys.

## Author Contributions

DB wrote the main manuscript text and prepared figures. All authors reviewed the manuscript.

## Additional Information

### Competing Interests Statement

The authors declare no competing interests.

## Supplementary Information Figure Caption

**Fig S1:**
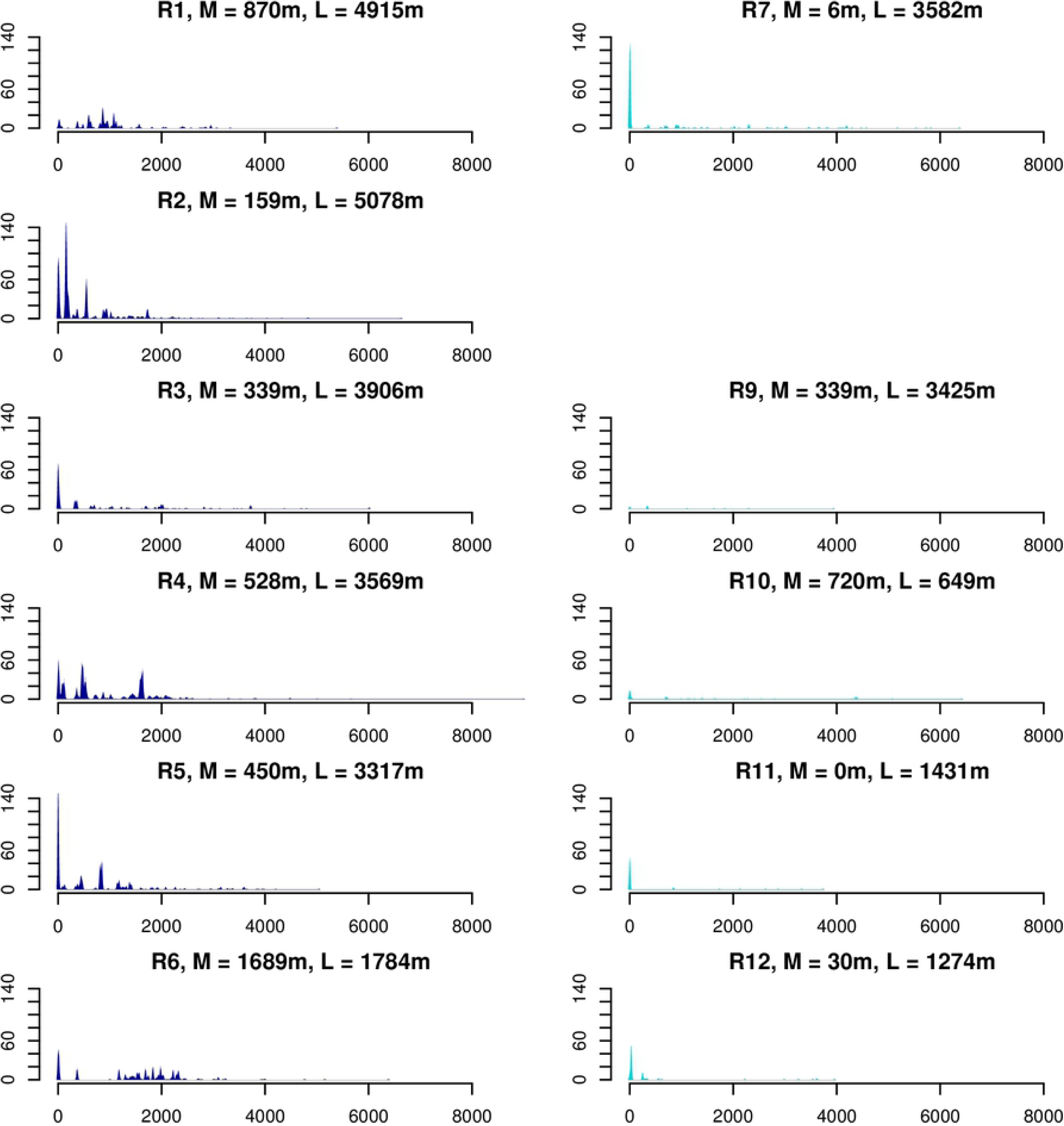
Raw distances between traps and fields. Minimal dispersal distances are on x-axis. Seeds number are on y-axis. Dark blue graphs correspond to two-lane roads and light blue graphs to one-lane roads. M stands for median as “the median minimal distances between trap-sites on the road and assigned fields”. L stands for length as “the total length of the road”.

